# Ocular TRPV1 deficiency protects from dry eye-induced corneal nerve damage

**DOI:** 10.1101/2023.08.21.554143

**Authors:** Manuela Pizzano, Alexia Vereertbrugghen, Agostina Cernutto, Florencia Sabbione, Irene A Keitelman, Carolina M Shiromizu, Douglas Vera Aguilar, Federico Fuentes, Mirta N Giordano, Analía S Trevani, Jeremías G Galletti

## Abstract

**Background:** Corneal nerve damage causes the most clinically significant symptoms in dry eye disease (DED) yet its pathophysiology remains poorly understood. Transient receptor potential vanilloid-1 (TRPV1) channels abound in corneal nerve fibers and respond to inflammation-derived ligands, which increase in DED. TRPV1 overactivation promotes axonal degeneration in vitro but whether it contributes to corneal neuropathy is unknown. Therefore, here we explored the role of TRPV1 in DED-associated corneal nerve damage.

**Methods:** Surgical DED was induced in TRPV1-deficient (TRPV1KO) and wild-type (wt) mice. Corneal nerve function was measured on days 0, 5, and 10 by mechanical and capsaicin sensitivity and eye-closing ratio as an indicator of non-evoked pain. Nerve and epithelial morphology was evaluated by confocal microscopy of corneal wholemounts. Pharmacological TRPV1 inhibition in wild-type mice was also evaluated.

**Results:** wt and TRPV1KO mice developed comparable ocular desiccation and corneal epithelial damage. Contrasting with wt mice, corneal mechanosensitivity in TRPV1KO mice did not decrease with disease progression. Capsaicin sensitivity increased in wt mice with DED, and consistently, wt but not TRPV1KO mice with DED displayed signs of non-evoked pain. Wt mice with DED exhibited nerve degeneration throughout the corneal epithelium whereas TRPV1KO mice only developed a reduction in the most superficial nerve endings that failed to propagate to the deeper subbasal corneal nerves. Pharmacological blockade of ocular TRPV1 activity reproduced these findings in wt mice with DED. Although TRPV1KO mice with DED had fewer pathogenic Th1 and Th17 CD4+ T cells in the lymph nodes, conjunctival immune infiltration was comparable between strains. Moreover, CD4+ T cells from wt and TRPV1KO mice with DED were equally pathogenic when transferred into T cell-deficient mice, confirming that TRPV1 activity in T cells is not involved in corneal neuropathy.

**Conclusions:** Although ocular desiccation is sufficient to trigger superficial corneal nerve damage in DED, proximal propagation of axonal degeneration requires TRPV1 signaling. Conversely, local inflammation sensitizes ocular TRPV1 channels, which are also involved in ocular pain, a key symptom of the disease. Thus, our findings suggest that ocular TRPV1 overactivation is a driving force in DED-associated corneal neuropathy and a potential therapeutic target.

**Graphical abstract:** 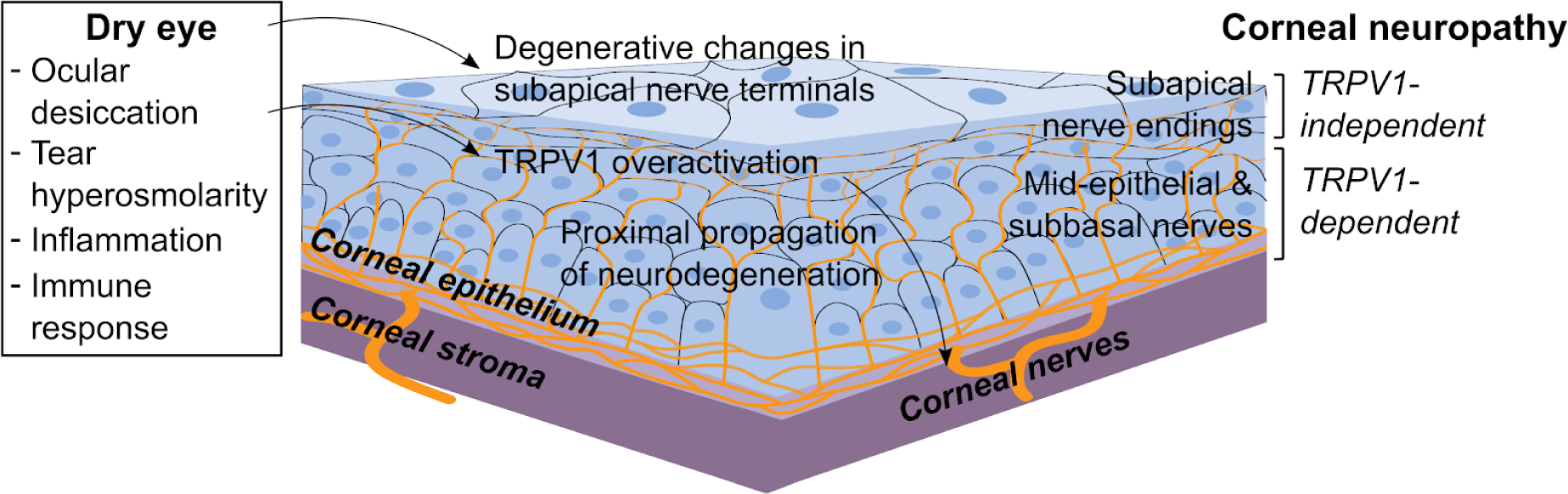

## Introduction

The cornea is the eye’s most powerful refractive lens but it can only fulfill its role when it is kept wet, smooth, and transparent(1). At the same time, the cornea is continuously exposed to rapidly changing environmental conditions that pose a threat to its integrity, and with it, to proper sight. To cope with this challenge, the cornea is sheltered within the ocular surface(2–4) and is endowed with abundant innervation that reaches peak density in its anteriormost layer, the corneal epithelium(5,6). The intraepithelial corneal nerve fibers detect fluctuations in moisture, temperature, and osmolarity and provide sensory input to the protective neural mechanisms that control blinking and tearing(7). Thus, corneal innervation plays an important role in corneal function and protection(7).

Dry eye disease (DED) is a multifactorial and increasingly prevalent ocular surface disorder typified by a dysfunctional tear film, which ultimately leads to ocular discomfort and pain, corneal damage, and impaired sight(8). Initially, desiccation causes tear hyperosmolarity and epithelial cell injury, both of which elicit an inflammatory response that aggravates tissue damage, thus creating a self-perpetuating vicious cycle(9). In addition to the ocular surface epithelium, corneal nerves are also affected in DED(7). Human studies show that there is a decrease in corneal nerve density, reduced fiber thickness, increased beading, tortuosity and sprouting, and signs of abnormal regeneration(10–19), and animal models of the disease reproduce these findings(20–25). Corneal nerve dysfunction in and of itself also reinforces DED(7,12,26). But contrasting our current understanding of how the corneal epithelium is affected by the disease, little is known about the pathophysiology of corneal neuropathy in DED.

Corneal innervation comprises a heterogeneous population of fibers that can be broadly classified by their functional and biochemical properties into three types: mechano-nociceptors, cold thermoreceptors, and polymodal nociceptors(7,26–28). The first two respond to pressure or temperature changes whereas polymodal nociceptors react to diverse noxious stimuli. Corneal polymodal nociceptor fibers typically express transient receptor potential vanilloid-1 (TRPV1) channels(7,12,29), which are activated by heat, low pH, hyperosmolarity, and various endogenous ligands that are increased in the context of inflammation and cell damage(30). Thus, TRPV1 signaling is critical to nociception and pain perception(31). In the skin, TRPV1 activation leads to proinflammatory neuropeptide release from local nerve fibers in a process known as cutaneous neurogenic inflammation(32). Consistently, we have previously shown that TRPV1 activation in the ocular surface also promotes substance P release with the subsequent neurogenic proinflammatory effects(33). On the other hand, TRPV1 overstimulation leads to neuronal and non-neuronal cell death in vitro(34–38) and corneal TRPV1 channel activity reportedly increases in the context of DED(39–43). However, whether TRPV1 signaling plays a role in the development of DED-associated corneal neuropathy remains unexplored.

Considering that a proinflammatory ocular surface milieu favors TRPV1 activation and that overstimulation of TRPV1 channels may have deleterious effects on cell membrane integrity and function according to in vitro models(34,35), we hypothesized that excessive TRPV1 signaling contributes to corneal nerve damage in DED. To this aim, we evaluated corneal neuropathy development in TRPV1-deficient (TRPV1KO) mice and also assessed the effect of pharmacological TRPV1 blockade on the corneal nerves of wild-type mice with DED. Altogether our results show that ocular TRPV1 signaling is required for corneal nerves to be affected in DED and thus have considerable therapeutic impact.

## Methods

### Mice

C57BL/6 (C57BL/6NCrl) mice were originally obtained from Charles River Laboratories (Wilmington, MA, USA). TRPV1KO (B6.129X1-*Trpv1^tm1Jul^*/J, JAX stock #003770) and recombination activating gene 1-deficient (RAG1KO, B6.129S7-*Rag1^tm1Mom^*/J, JAX stock #002216) mice were purchased from The Jackson Laboratory (Bar Harbor, ME, USA). Mice were bred and maintained at the Institute of Experimental Medicine’s animal facility. All mice were 6-8 weeks old at the beginning of the experiments and both male and female mice were included. All protocols were approved by the Institute of Experimental Medicine animal ethics committee and adhered to the Association for Research in Vision and Ophthalmology Statement for the Use of Animals in Ophthalmic and Vision Research.

### Reagents, antibodies, and cell cultures

All chemical and biological reagents were from Sigma-Aldrich (Buenos Aires, Argentina) unless otherwise specified. Fluorochrome-tagged antibodies were from Biolegend (San Diego, CA, USA) unless otherwise specified. All cell cultures were done in RPMI-1640 medium supplemented with 10% fetal calf serum, 10 mM glutamine, 100 U/ml penicillin, 100 μg/ml streptomycin, and 5x10-5 M 2-mercaptoethanol in a humidified incubator with 5% CO2 at 37°C.

### Lacrimal gland excision surgery

Mice were anesthetized by i.p. injection of ketamine (100 mg/kg) and xylazine (10 mg/kg). Excision surgery comprised four steps: first, a 3-mm-long incision was made along the middle third of the line joining the lateral canthus of the ear and the pinna; second, the superior pole of the extraorbital lacrimal gland was exposed by incising the fibrous capsule that surrounds it; third, the lacrimal gland was pulled out gently and excised taking special care not to damage the blood vessels next to its inferior pole; fourth, the skin was closed using 6-0 nylon thread. The glands from both sides were excised sequentially. Sham surgery consisted of only steps 1 and 4. In all cases, a single dose of 10 mg/kg diclofenac sodium was injected s.c. in the scruff for postoperative analgesia and ciprofloxacin ointment as applied over the wound once the surgery was completed. The eyes were protected from desiccation with sodium hyaluronate 0.4% (Dropstar LC, Laboratorio Poen, Argentina) until the mice recovered from anesthesia.

### Assessment of tear production

Tear production was measured by inserting a 1 mm-wide phenol-red impregnated filter paper strip in the inferior conjunctival fornix adjacent to the lateral canthus, where it was held in place for 60 seconds while restraining the mouse gently and allowing for normal blinking. The wetted length of the right eye of each mouse was measured and used as a data point.

### Assessment of corneal epithelial barrier function

Corneal fluorescein uptake was measured as previously described(44). In brief, 0.5 μl of dextran-fluorescein isothiocyanate (average mol wt 3,000-5,000, 10 mg/ml in PBS) was applied to each eye and then the mouse was returned to its cage. After 3 min, a 10-20 second-long video of each eye under blue light was captured with the aid of a fluorescence-adapted dissection microscope (NightSea SFA-RB). For analysis, a masked observer exported a representative video frame as an image and selected the corneal area suitable for analysis, excluding reflections and other artifacts, using ImageJ software. The mean fluorescence intensity of the resulting region of interest was calculated after background subtraction (50-pixel rolling ball radius), and the average of both eyes was used for analysis.

### Assessment of corneal mechanical sensitivity

Mechanical thresholds were determined using a mouse-adapted version of Cochet-Bonnet esthesiometry(7,34). Nylon 6-0 monofilament was cut into segments of varying lengths (1.0 to 5.5 cm in 0.5 cm steps). With the mouse held firmly in one hand, the cornea was touched six times with each filament, starting with the longest segment. A positive response was defined as blinking and retraction of the eye in reaction to at least three of the six tries. The longest segment yielding a positive response was used as the sensitivity threshold, and the average of both eyes was used for analysis. Corneal sensitivity was measured in the morning (8-11 AM) before any other experimental manipulation.

### Eye-closing ratio

One mouse at a time was placed on an elevated platform and allowed to habituate for 2 min. Then, a >1-min-long video was recorded with a camera placed at the same height. For analysis, a masked observer selected snapshots in which each eye was clearly visible. The distance between canthi (x) and between the upper and lower lids (y) was measured using ImageJ and then the corresponding eye-closing ratio was calculated as y/x. At least two snapshots per eye were analyzed, and then the results from both eyes were averaged to obtain one data point per mouse.

### Capsaicin sensitivity

Eye wiping behavior was measured in response to 100 μM capsaicin. Immediately after applying 5 μl solution onto each eye, the mouse was placed in a separate cage and recorded with a camera placed above for at least 30 seconds. For analysis, a masked observer counted the number of eye wipes during the first 30 seconds using a slow playback speed.

### Spleen and lymph node cells

Cervical, axillary, and inguinal lymph nodes were harvested after euthanasia and rendered into a cell suspension by mechanical dissociation through nylon mesh. For splenocyte suspensions, red blood cells were lysed with ammonium chloride-potassium buffer.

### Intracellular cytokine staining

Cells were stimulated in U-bottom 96-well plates (0.5x10^6^ cells/0.2 ml media/well) for 5 h with 50 ng/ml phorbol myristate acetate, 1 µg/ml ionomycin, and 10 µg/ml brefeldin A. DNAse (1 U/ml) was added 15 min before the stimulation period ended. The cells were then washed, stained with a fixable viability dye (#L34975, Thermo Fischer Scientific, Buenos Aires, Argentina), washed, stained with CD4 (CD4 FITC #100406, Biolegend), and then fixed, permeabilized, and stained for intracellular cytokines (IFN-γ PE #505808, IL-4 BV421 #504120, IL-17A PE-Cy7 #506922, Biolegend) with the Cyto-Fast™ Fix/Perm Buffer Set (#426803, BioLegend) as per the manufacturer’s instructions.

### Preparation and flow cytometry analysis of conjunctival cell suspensions

Conjunctival tissue was minced into fragments with the aid of scissors, incubated in collagenase 1 mg/ml in PBS at 37° C with gentle shaking for 30 min, then DNAse 2 U/ml was added and the tissue samples were digested for another 15 min. Digestion was stopped by adding 2 mM EDTA and 10% fetal calf serum and the suspension was washed and filtered for staining. Cell suspensions were first stained with a fixable viability dye (#L34975, Thermo Fischer Scientific, Buenos Aires, Argentina), washed, Fc-receptor blocked, stained for surface markers (CD45 APC #103112, CD4 FITC #100406, Ly6G PE-Cy7 #127618, Biolegend), and then fixed in 1% paraformaldehyde in PBS. The entire cell suspension resulting from one mouse was stained and acquired for analysis as one independent sample. First, singlets were gated based on forward scatter height versus area, then gated on side scatter height versus side scatter area, then gated on viability dye-excluding events (viable cells), and finally on CD45^+^ CD4^+^ or CD45^+^ Ly6G^+^ events.

### Collection of eye tissue

After euthanasia, the conjunctival tissue of each eye was excised as two strips (superior and inferior) under a dissection microscope and collected in ice-cold RPMI media without serum. Immediately after, enucleation was performed by gently proptosing the eye globe and cutting the optic nerve with curved scissors. The two eyes of each mouse were collected in ice-cold fixative solution. Mice were euthanized one at a time so that all ocular tissue was collected within 5 minutes of the time of death to ensure adequate corneal nerve preservation(45).

### Corneal immunostaining

Eyes were processed as described by Tadvalkar et al(45). In brief, eyes were fixed in a pre-chilled formaldehyde-containing buffer for 75 min, washed, and stored in methanol at -20° C until processed for staining. Then, the fixed corneas were cut from the back of the eye under a dissection microscope, permeabilized with a graded methanol-Triton X-100 series, blocked overnight with 1% BSA and 1% goat serum in PBS, and stained overnight with Alexa 488-conjugated Alexa Fluor® 488 anti-tubulin β3 and Alexa Fluor® 647 anti-mouse/human CD324 (E-cadherin) antibodies (#801203 and #147308, BioLegend). Each batch of anti-tubulin β3 antibody was titrated before use to minimize background staining, usually resulting in 0.5-0.7 μl antibody/200 μl buffer/cornea (2.5-3.5 μg/ml) as optimal. The stained corneas were washed three times for 60 min in PBS-Tween 0.02%, counterstained with 1 ug/ml DAPI, mounted flat with the aid of relaxing cuts in Aqua-Poly/Mount (PolySciences), and stored at 4° C until imaged.

### Confocal laser scanning microscopy acquisition

Image acquisition was performed with a FluoView FV1000 confocal microscope (Olympus, Tokyo, Japan) equipped with Plapon 60X/1.42 and UPlanSapo 20X/0.75 objectives. Z stacks (0.5-μm step size) spanning the entire corneal epithelium (approximately 30 μm) were obtained at two opposite located at 600 μm from the center (defined as the center of the nerve whorl or as the center of the disorganized area in those samples with highly disrupted nerve whorls). Corneal nerve analysis was performed at three different levels within the corneal epithelium. For subapical nerve terminals, the first section located entirely beneath the apical epithelial squames (1-1.5 μm deep, usually the third or fourth) was selected. Then, the image was thresholded after background subtraction (10-pixel rolling ball radius), and the percentage area occupied by nerve endings was determined by the corresponding ImageJ function. For mid-epithelial nerve terminals, the mid-section (60X) between the apical- and basal-most sections from each stack was chosen. Then, the number of nerve endings was assessed after background subtraction (10-pixel rolling ball radius) by a masked observer using the Cell Counter ImageJ function. Data are shown as the number of terminals/60X field (423.94 µm^2^ area). To analyze the complexity of the subbasal epithelial nerves, the Sholl plugin in ImageJ software was used. In brief, a maximum intensity projection of the 10 sections encompassing the corneal subbasal nerve mat was created, then the background was subtracted (50-pixel rolling ball radius), and the image was thresholded. Finally, 10 concentric circles with a 10-µm radius step size were traced at the center of the final image and the resulting sum of intersections of tubulin β3^+^ nerves for each concentric circle was calculated using the software and used for analysis(33). Corneal epithelial nuclei were counted using the built-in neural network model for fluorescent nuclei of the ImageJ plugin StarDist, a cell/nuclei detection method(46). The nuclei count in 30 sections spaced 1 µm apart starting at the basalmost epithelial section and progressing apically were added to obtain one data point.

### RT-qPCR analysis of trigeminal gene expression

Euthanized mice were transcardially perfused with phosphate-buffered saline. Both trigeminal ganglia from each mouse were isolated under a dissection microscope, collected in ice-cold Trizol, and stored at -70 °C as one sample. RNA was extracted with Direct-zol RNA MiniPrep columns (Zymo Research, USA) as per the manufacturer’s instructions and reverse transcription was performed as previously described(33). Real-time quantitative PCR was performed with 50 ng cDNA and SsoFast™ EvaGreen® Supermix (Bio-Rad, USA) and primers in a final reaction volume of 20 μL. Primers were designed using the Primer3 software, purchased from Ruralex-Fagos (Buenos Aires, Argentina), and used at 400 nM: glyceraldehyde 3-phosphate dehydrogenase Fw 5′ CTCCCACTCTTCCACCTTCG 3′, Rv 5′ CCACCACCCTGTTGCTGTAG 3′; TRPV1 Fw 5′ CATCTTCACCACGGCTGCTTAC 3′, Rv 5′ CAGACAGGATCTCTCCAGTGAC 3′; transient receptor potential melastatin-8 (TRPM8) Fw 5′ GTTGGACCTTGCCAGTGATGAG 3′, Rv 5′ CCATTCTCCAGAAAGAGGCGGA 3′; Piezo2 Fw 5′ GCACTCTACCTCAGGAAGACTG 3′, Rv 5′ CAAAGCTGTGCCACCAGGTTCT 3′; activating transcription factor-3 Fw 5′ GAAGATGAGAGGAAAAGGAGGCG 3′, Rv 5′ GCTCAGCATTCACACTCTCCAG 3′; tumor necrosis factor Fw 5′ CTTCTCATTCCTGCTTGTGG 3′, Rv 5′ GGGAACTTCTCATCCCTTTG 3′. The reaction was performed in a CFX Connect Real-Time PCR Detection System (Bio-Rad, USA). The glyceraldehyde 3-phosphate dehydrogenase gene was used for normalization of the results for each sample and then the fold-change in specific mRNA levels between groups was calculated by using the 2^−ΔΔCt^ method.

### Isolation and adoptive transfer of CD4+ T cells

CD4^+^ cells were isolated from pooled splenocytes and lymph node cells with the aid of magnetic beads (MojoSort™ Mouse CD4^+^ T Cell Isolation Kit, BioLegend #480033) as per the manufacturer’s instructions. Cell purity was >95% as assessed by flow cytometry (CD4 staining). For adoptive transfer experiments, cells were resuspended in PBS and 1x10^6^ cells/0.5 ml were injected intraperitoneally into each RAG1KO recipient mouse.

### Statistical analysis

Student’s t-test and one- or two-way analysis of variance (ANOVA) with Bonferroni or Dunnett’s posthoc tests were used to compare the means of two or more samples, respectively. Significance was set at p<0.05 and two-tailed tests were used in all experiments. Calculations were performed using GraphPad Prism version 7 software (GraphPad Software, La Jolla, Ca, USA).

## Results

### 1 TRPV1 deficiency does not impact tear production or corneal epithelial integrity in the context of DED

Among other stimuli, TRPV1 channels are activated by endogenous lipid-derived inflammatory mediators and hyperosmolar conditions(47), both of which are found in the DED-affected cornea. In line with this, we have previously shown that tear hyperosmolarity, the natural consequence of increased desiccation in DED, enhances TRPV1 signaling in the cornea(41). We also found that TRPV1 activation in the context of a corneal alkali burn leads to neurogenic inflammation(33), and others reported that TRPV1 overstimulation may lead to cell membrane disruption in vitro(34,35). Herein we hypothesized that TRPV1 signaling could be involved in DED-associated corneal nerve degeneration. To test this, we compared the development of DED in wild-type and TRPV1KO mice (Figure 1A) using a previously characterized surgical model of the disease (Figure 1B) that involves the bilateral excision of the extraorbital lacrimal gland (48–50). This model corresponds to an aqueous-deficient form of DED because the mice exhibit a steady, marked reduction in basal tear production.

**Figure 1.**
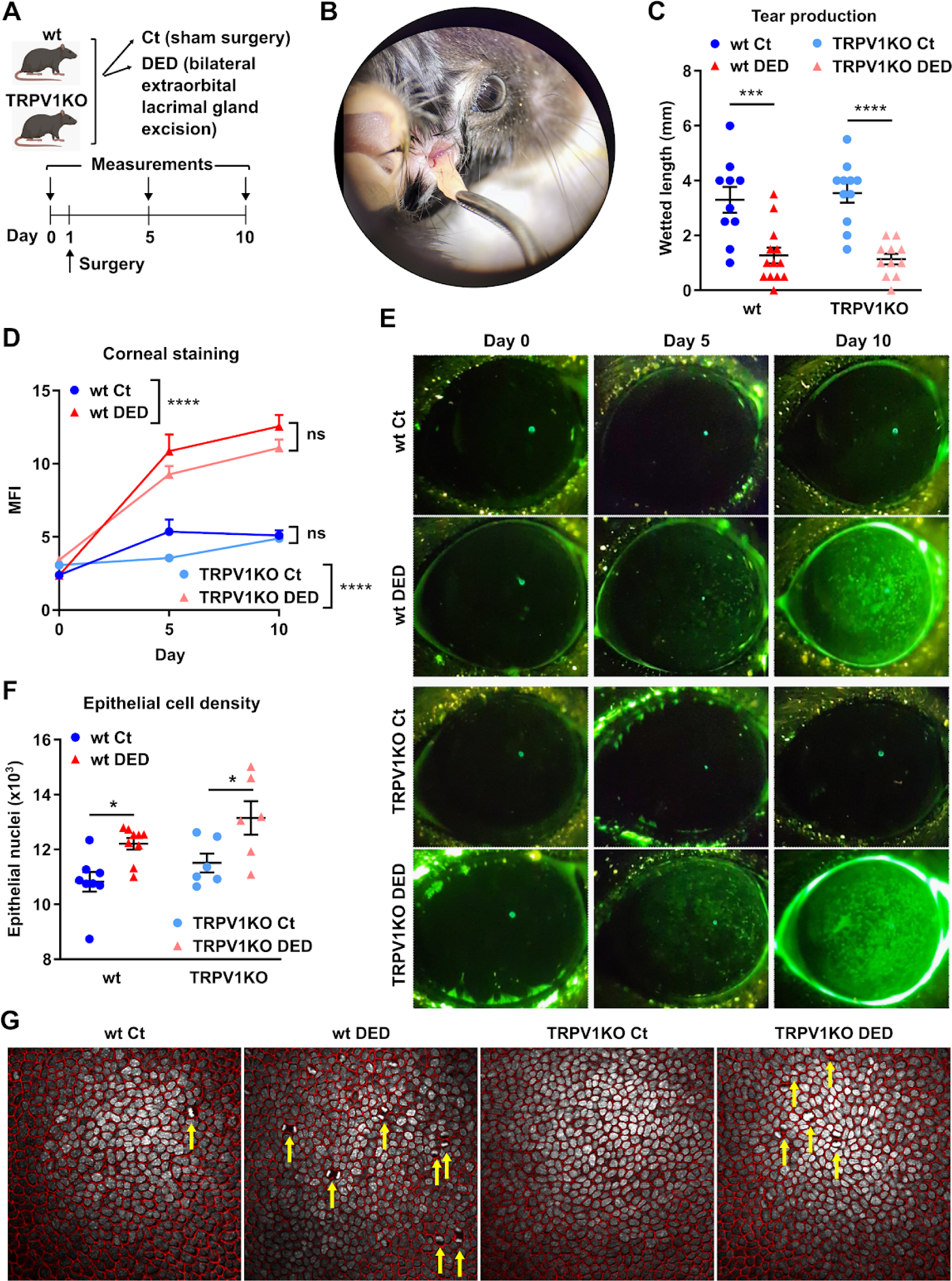
TRPV1 deficiency does not impact tear production or corneal epithelial integrity in the context of dry eye. A) Dry eye disease (DED) was surgically induced in wild-type (wt) or transient receptor potential vanilloid-1-deficient (TRPV1KO) mice through bilateral excision of the extraorbital lacrimal gland. Sham-operated animals were included as controls (Ct). A) Experimental design and B) surgical technique. C) Tear production on day 5 as measured by phenol red-paper wetting length. D) Cumulative data and E) representative micrographs of corneal dextran-fluorescein uptake in Ct and DED mice from both strains. F) Total nuclei count in 30 sections spanning 30 μm of corneal epithelium. G) Representative micrographs of basal corneal epithelium stained with e-cadherin (red) and DAPI (white). Mitotic figures are highlighted with yellow arrows. All experiments were performed twice or more with 6 mice/group/experiment. For all experiments, mean±standard error of measurement is shown. To compare means, two-way ANOVA was used for C (strain and treatment), D (group and time), and F (strain and treatment). Sidak’s posthoc test was applied in all cases. * indicates p<0.05, *** indicates p<0.001, **** indicates p<0.0001, and ns indicates not significant.

Although basal tearing is regulated by TRPM8 channels expressed in corneal cold thermoreceptors(51), there is evidence of TRPV1-mediated sensitization of these cooling-responsive fibers in DED(52). Therefore, we first verified that ocular surface desiccation was comparable in both strains to rule out a confounding effect of TRPV1 deficiency on this aspect of the model. To this aim, we assessed tear production levels in wild-type and TRPV1KO mice after sham and lacrimal gland excision surgery (Figure 1C). Tear production in sham-operated mice of both strains did not differ (p=0.99), as reported elsewhere(39). Consistently, lacrimal gland excision reduced tear production in wt and TRPV1KO mice to a comparable extent (62% vs 68%, p=0.99). Thus, mouse strain had no effect on the tear production observed in both treatment groups (2-way ANOVA: factors strain p=0.87, treatment p<0.0001). Next, we evaluated the development of corneal epitheliopathy as a hallmark of DED. Corneal epithelial damage can be graded by assessing the uptake of a fluorescent probe (Figure 1D) as an indicator of corneal epithelial barrier function(53). As shown in Figure 1E, corneal staining increased comparably in wt and TRPV1KO mice with DED (p<0.0001 compared with same-strain control mice) and there was no significant difference between strains with the same treatment at any time point (p>0.05). Finally, we examined the epithelial morphology in corneal wholemounts because DED increases cell turnover, which is accompanied by a decrease in epithelial cell size(54,55). As shown in Figure 1F, both wild-type and TRPV1KO mice with DED had more epithelial cells within the same volume of analyzed corneal epithelium (treatment factor p=0.0005), and in line with this, more mitotic figures were evident in the basal epithelial layers of mice with DED from both strains (Figure 1G). Of note, TRPV1KO mice exhibited slightly higher nuclei counts than same-treatment wild-type animals (strain factor p=0.04). Both findings point towards comparably increased cell proliferation in response to desiccation in both strains, and altogether the data shows that DED-associated changes in tear production and in the corneal epithelium were comparable in wild-type and TRPV1KO mice.

### 2 TRPV1 deficiency prevents DED-induced impairment of corneal nerve function

Having established that wild-type and TRPV1KO mice with DED develop comparable ocular desiccation and corneal epitheliopathy, we then assessed the extent of DED-induced corneal neuropathy in both strains. We first measured corneal mechanical sensitivity using a mouse-adapted version of Cochet-Bonnet esthesiometry, a widely-used technique to measure corneal nerve function in the clinic that relies on the fact that longer filaments exert less pressure upon corneal contact (Figure 2A). Unexpectedly, TRPV1KO mice had a significantly lower pressure threshold (4.45±0.357 vs 3.97±0.279 cm, p<0.0001) at baseline, indicating greater sensitivity to mechanical stimulation than wild-type mice. As previously reported(21), wild-type mice with DED showed a progressive decline in mechanosensitivity on days 5 and 10 compared to wild-type controls (Figure 2A). By contrast, TRPV1KO mice with DED did not experience a change in their corneal mechanosensitivity thresholds. This finding indicates that wild-type mice experienced more DED-associated corneal nerve dysfunction than TRPV1KO mice. Considering the different starting thresholds between strains, relativization of measurements to the same-strain baseline better shows that the DED-associated change only occurred in wild-type mice (Figure 2B). We also measured corneal sensitivity to capsaicin, a well-characterized agonist of TRPV1 channels that elicits ocular pain and nocifensive behavior in mice (Supplementary Videos 1 and 2). Over the 10-day span of the experiment, capsaicin instillation elicited a constant number of wipes in sham-operated wild-type mice whereas this nocifensive response progressively increased in wild-type mice with DED (Figure 2C). This finding indicates that DED brings about sensitization of TRPV1-expressing corneal fibers, suggesting that TRPV1 signaling increases with the inflammatory context during disease progression. As expected, capsaicin did not evoke a wiping response in TRPV1 mice (data not shown). Finally, we analyzed the eye-closing ratio (Figure 2D and 2E), which is part of the validated Mouse Grimace Scale for quantifying the spontaneous subjective pain experience in mice(56,57). Wild-type mice with DED showed signs of spontaneous pain while TRPV1KO mice with DED did not exhibit differences in the eye-closing ratio compared with sham-operated mice of both strains. This result indicates that DED brings about TRPV1-dependent nociception that ultimately results in pain sensation. Altogether these findings show that DED causes TRPV1 sensitization and activation, and more importantly, that these events are accompanied by corneal nerve dysfunction.

**Figure 2.**
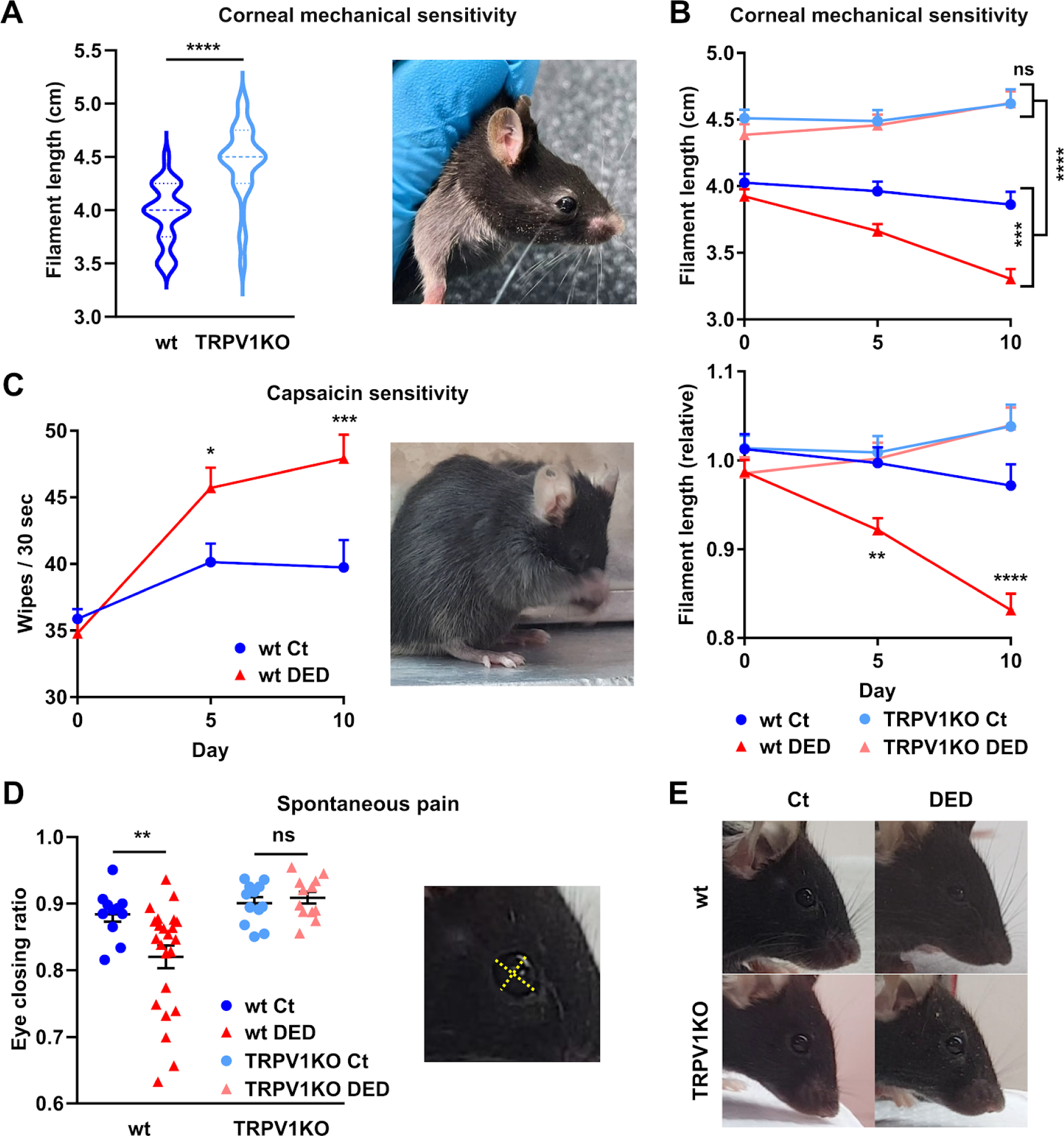
TRPV1 deficiency prevents dry eye-induced impairment of corneal nerve function. A) Corneal mechanosensitivity was measured in wild-type (wt) or transient receptor potential vanilloid-1-deficient (TRPV1KO) mice before each experiment. Pooled data (left) from 43 wt and 47 TRPV1KO mice and representative image (right) showing the modified Cochet-Bonnet esthesiometry technique. B) Corneal mechanosensitivity in wt and TRPV1KO mice with surgically induced dry eye disease (DED) or sham surgery (Ct). Measurements are shown as absolute data (top) or relative to the same-strain baseline. C) Capsaicin sensitivity as measured by the number of eye wipes elicited over 30 seconds following the TRPV1 agonist instillation on both eyes. Cumulative data (left) and representative example (right) of the nocifensive behavior counted as a positive response. D) Eye-closing ratio measurements in Ct and DED mice from both strains. Cumulative data (left) and analysis technique (right). E) Representative images of mice from both treatment groups and strains depicting the use of the eye-closing ratio as an indicator of spontaneous ocular pain. All experiments were performed twice or more with 6 mice/group/experiment. For all experiments, mean±standard error of measurement is shown. To compare means, Student’s t test was applied in A, two-way ANOVA with Sidak’s posthoc test was used for B and C (group and time) and D (strain and treatment). * indicates p<0.05, ** indicates p<0.01, *** indicates p<0.001, **** indicates p<0.0001, and ns indicates not significant.

### 3 TRPV1 deficiency reduces DED-induced changes in corneal nerve structure

As we observed no DED-associated corneal nerve dysfunction in TRPV1KO mice, we examined nerve morphology by confocal microscopy of corneal whole mounts. The morphology of the intraepithelial corneal nerves was qualitatively comparable between sham-treated wild-type and TRPV1KO mice, with the typical whorl pattern of the subbasal plexus. Axial reconstructions of the central intraepithelial corneal nerves (Figure 3A) confirmed the conserved morphology in sham-treated TRPV1KO mice. Compared to sham-treated mice, reconstructions from wild-type mice with DED showed focal reductions of innervation that were more evident at the mid-epithelial level, where the nerve fibers run perpendicularly towards the corneal surface (Figure 3A and 3B). En face imaging at three distinct depths within the corneal epithelial layer (Figure 3C) allowed for quantification and comparison between groups (Figure 3D, E, and F). Baseline innervation at each level differed between strains: TRPV1KO mice have higher subapical nerve-occupied area but lower mid-epithelial nerve density and subbasal nerve complexity. To account for this source of variation and better reflect DED-induced changes, data was relativized to sham-treated mice of the same strain. As shown in Figure 3D, DED comparably decreased the total area occupied by subapical nerve endings beneath the apicalmost layer of corneal epithelial cells in wild-type and TRPV1KO mice. By contrast, wild-type mice with DED experienced a drop in the mid-epithelial and subbasal nerve density whereas TRPV1KO mice with DED did not exhibit such a change (Figure 3E and 3F). Altogether these results show that the pattern and extent of DED-associated morphological nerve changes in TRPV1KO mice is not the same as in wild-type mice. Despite baseline differences, DED induces a comparable loss of the most exposed subapical endings while the deeper portion of the intraepithelial nerves remain relatively unchanged.

**Figure 3.**
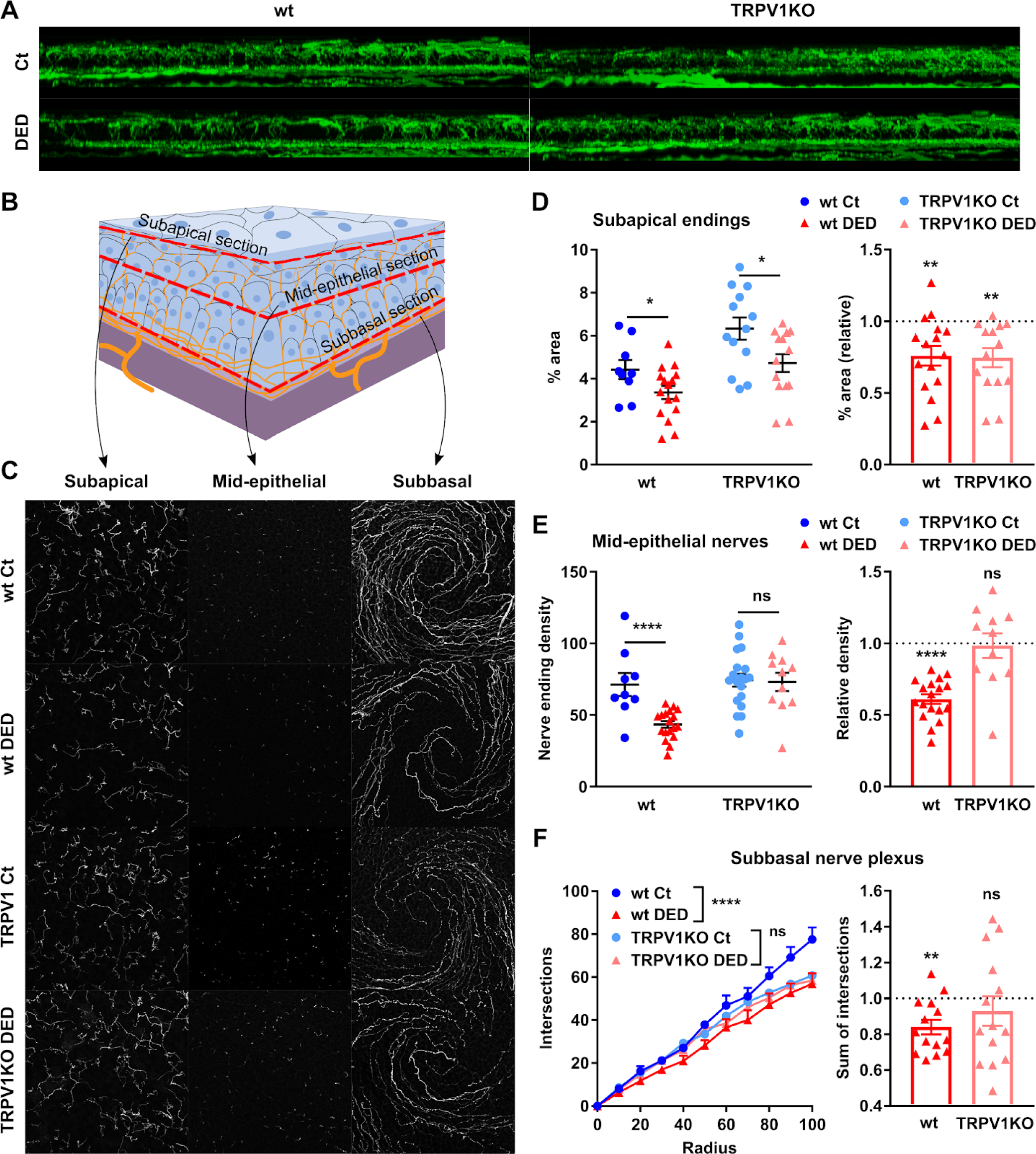
TRPV1 deficiency reduces DED-induced changes in corneal nerve structure. Corneal wholemounts from wild-type (wt) or transient receptor potential vanilloid-1-deficient (TRPV1KO) mice with surgically induced dry eye disease (DED) or sham surgery (Ct) were stained with Alexa Fluor® 488 anti-tubulin β3 (shown in green or white). A) Axial reconstructions of the corneal nerves spanning the entire epithelial layer and the anterior stroma. Representative images from Ct and DED mice from both strains. B) Schematic of the levels (subapical, mid-epithelial, and subbasal) at which corneal nerve morphology was analyzed and C) representative micrographs for each level from all groups. D) Quantification (% area) of the most superficial nerve terminals imaged en face beneath the apical epithelial squames (subapical section). E) Density (count of nerve endings/field) of intraepithelial corneal nerves in cross-section at the mid-epithelial level as they run perpendicularly to the surface. For D and E, measurements are shown as absolute data from all groups (left) or the DED group from each strain relative to the same-strain Ct group (right). F) Quantification of corneal neural complexity at the subbasal level by Sholl analysis. Absolute number of intersections at each Sholl radius from all groups (left) or sum of intersections of the DED groups from both strains relative to the same-strain Ct group (right). All experiments were performed twice or more with 6 mice/group/experiment. For all experiments, mean±standard error of measurement is shown. To compare means, two-way ANOVA with Sidak’s posthoc test was used in the left panels of D and E (treatment and strain) and F (group and radius) whereas the one-sample Student’s t test was applied in the right panels of D, E, and F. * indicates p<0.05, ** indicates p<0.01, **** indicates p<0.0001, and ns indicates not significant.

### 4 TRPV1 deficiency reduces DED-induced changes in corneal nerve structure

Axonal degeneration in DED induces reactive changes in gene expression in the trigeminal ganglion, where the cell bodies of the cornea-innervating neurons are located(12,24,43). As reported by others(39,40), we observed a significant increase in *TRPV1* expression in the trigeminal ganglia of wild-type mice after 10 days of DED (Figure 4A, 1.04 ± 0.27 vs 1.96 ± 0.30-fold, p=0.04). As expected, *TRPV1* expression was not detectable in TRPV1-deficient mice (data not shown). The expression levels of *TRPM8* (transient receptor potential melastatin 8) and *Piezo2,* the two sensory channels typically associated with corneal cold-thermoreceptor and selective mechanonociceptor fibers, were also increased in wild-type DED mice (Figure 4B: *TRPM8* 1.00 ± 0.20 vs 2.47 ± 0.34-fold, p=0.001; Figure 4C: *Piezo2* 1.00 ± 0.30 vs 2.67 ± 0.49-fold, p=0.02). By contrast, TRPV1-deficient mice did not significantly modify their trigeminal expression of TRPM8 and Piezo2 in response to DED (Figure 4B: *TRPM8* 1.23 ± 0.27 vs 1.47 ± 0.23-fold, p=0.75; Figure 4C: *Piezo2* 1.23 ± 0.32 vs 0.93 ± 0.26-fold, p=0.49). Consistently, expression levels of activating transcription factor 3 (*ATF3*), a transcription factor that is rapidly upregulated in sensory neurons in response to axonal injury(58) increased in the setting of DED in wild-type mice (Figure 4D, 0.92 ± 0.24 vs 2.15 ± 0.38-fold, p=0.007) but not in TRPV1-deficient mice (0.72 ± 0.16 vs 1.14 ± 0.22-fold, p=0.47). However, expression of tumor necrosis alpha (Figure 4E) increased in both wild-type and TRPV1KO mice with DED (treatment factor: 0.001, strain factor: p=0.31), suggestive of neuroinflamatory changes in response to ocular surface disease. Altogether, these findings show that some reactive changes in trigeminal gene expression observed in wild-type mice with DED are attenuated or absent in TRPV1KO mice, which is consistent with the protective effect of TRPV1 deficiency on corneal neuropathy development.

**Figure 4.**
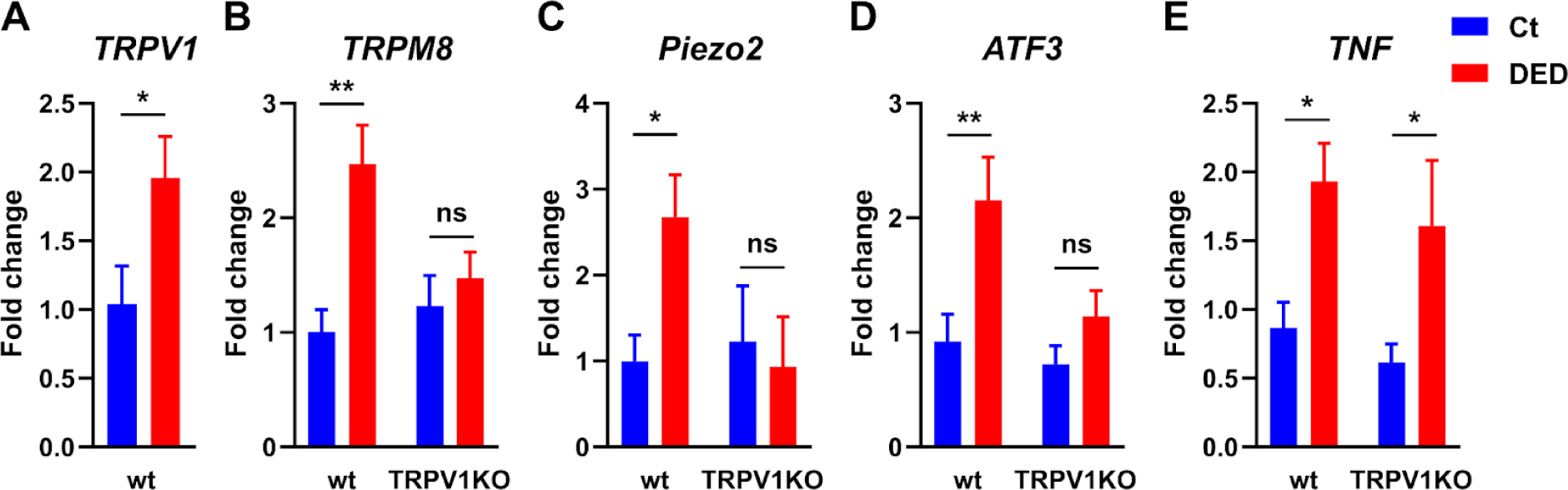
TRPV1 deficiency attenuates DED-induced gene expression changes in the trigeminal ganglion. Trigeminal ganglia from wild-type (wt) or transient receptor potential vanilloid-1-deficient (TRPV1KO) mice with surgically induced dry eye disease (DED) or sham surgery (Ct) were harvested on day 10 and assayed for mRNA expression levels of A) transient receptor potential vanilloid-1 (*TRPV1*), B) transient receptor potential melastatin-8 (*TRPM8*), C) *Piezo2*, D) activating transcription factor-3 (*ATF3*), and E) tumor necrosis factor (*TNF*) genes. Mean±standard error of measurement is shown, n=6 or more/group. To compare means, Student’s t test was applied in A and two-way ANOVA with Sidak’s posthoc test was used from B through E (treatment and strain). * indicates p<0.05, ** indicates p<0.01, **** indicates p<0.0001, and ns indicates not significant.

### 5 Pharmacological TRPV1 blockade in wild-type mice with DED prevents corneal neuropathy development

Since TRPV1KO mice exhibited some baseline differences in corneal nerve function (Figure 2A) and morphology (Figure 3D-E-F) that could confound the results, we tested the effect of pharmacological TRPV1 inhibition in wild-type mice with DED (Figure 5A). To this aim, we resorted to ocular instillation of the TRPV1 antagonist SB-366791 with a dosing scheme that has been previously validated in mice (4 times/day, 100 μg/ml)(33,41). We quantified the progression of corneal epitheliopathy and neuropathy using the same methodology as before. As assessed by corneal fluorescent probe uptake and in agreement with our findings in TRPV1KO mice, ocular TRPV1 blockade had no discernible effect (p=0.56) on the development of DED-induced corneal epitheliopathy in wild-type mice (Figure 5B and 5C). By contrast, topical TRPV1 inhibition ameliorated the corneal mechanical sensitivity drop associated with DED progression (Figure 5D, p=0.0018 for treatment). Although this finding is in line with the knockout-derived observations (Figure 3), TRPV1 inhibitor-treated mice did experience some loss in corneal mechanosensitivity after 10 days of DED (-11% from baseline, p=0.02), which was still significantly less than that of saline-treated mice at the same timepoint (-23% from baseline, p=0.03). As shown in Figure 4E, corneal nerve morphology changes in PBS-treated mice were similar to those previously observed in wild-type mice with DED (Figure 3A) whereas TRPV1 inhibitor-treated mice showed better preservation of the mid-epithelial and subbasal nerve segments, as did TRPV1KO mice with DED (Figure 3A). Quantification of corneal nerves revealed that TRPV1 antagonist instillation caused no statistically significant change in the subapical (most superficial) nerve endings (Figure 5F). By contrast, TRPV1 inhibition ameliorated DED-induced nerve fiber loss in the mid-epithelial vertical fibers (-39% vs -22%, p=0.01, Figure 5G) and in the deeper subbasal nerve plexus (-19% vs -6%, p<0.05, Figure 5H). Representative micrographs (Figure 5I) obtained at the three different levels within the corneal epithelium better depict the corneal nerve changes. These results confirmed the findings in TRPV1KO mice: ocular TRPV1 signaling is associated with corneal neuropathy but not corneal epitheliopathy development in DED.

**Figure 5.**
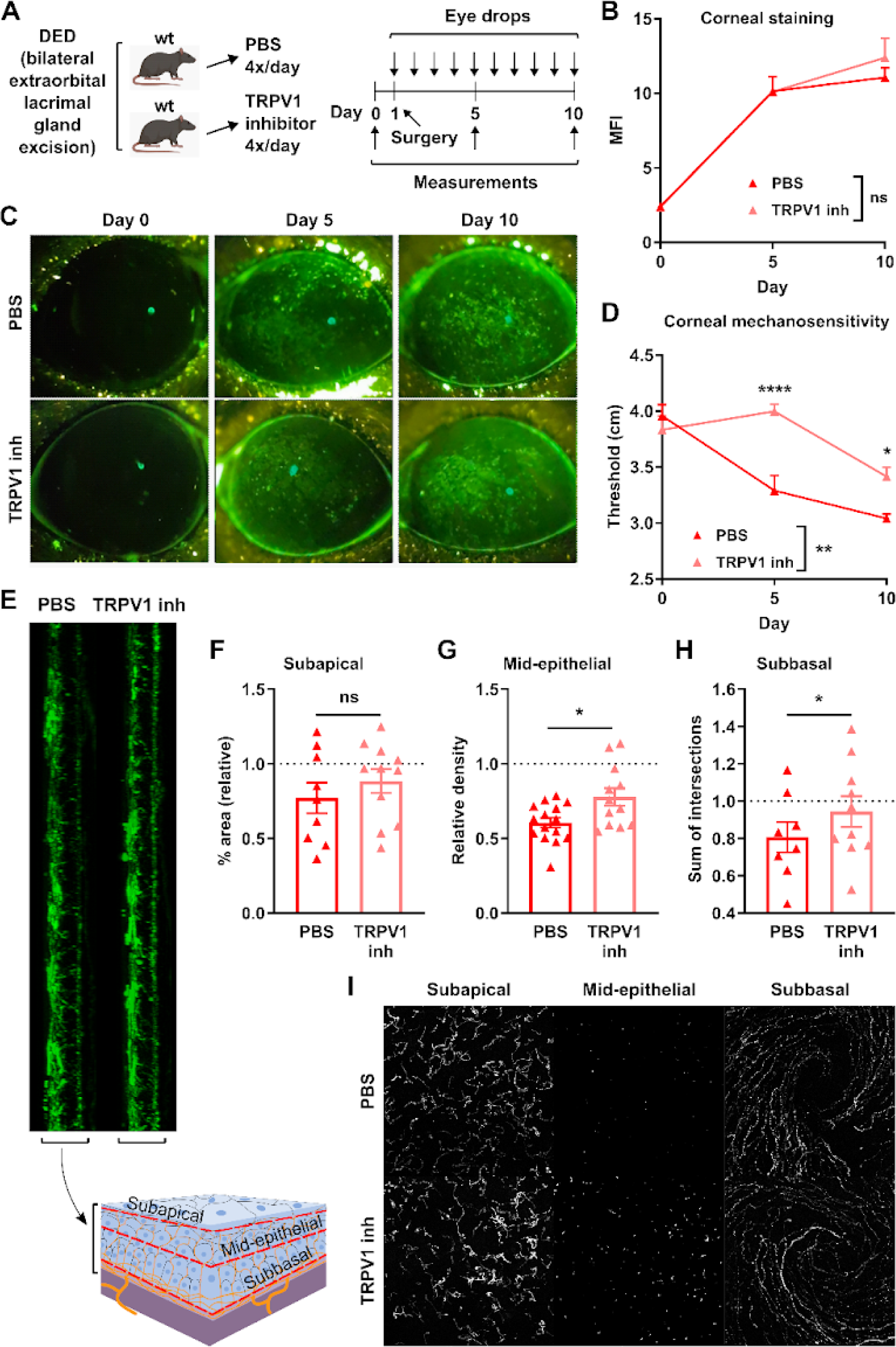
Pharmacological TRPV1 blockade in wild-type mice with DED prevents corneal neuropathy development. A) Dry eye disease (DED) was surgically induced in wild-type (wt) mice that after recovering from anesthesia started receiving eye drops (4 times/day) containing either phosphate-buffered saline (PBS) or 100 μg/ml SB-366791, a known transient receptor potential vanilloid-1 inhibitor (TRPV1 inh). B) Cumulative data and C) representative micrographs of corneal dextran-fluorescein uptake in PBS and TRPV1 inhibitor-treated DED mice. D) Corneal mechanosensitivity in PBS and TRPV1 inhibitor-treated DED mice. E) Axial reconstructions (top) of the central corneal nerves spanning the entire epithelial layer and the anterior stroma (tubulin β3 staining), and schematic (bottom) of the levels (subapical, mid-epithelial, and subbasal) at which corneal nerve morphology was analyzed. F) Quantification (% area) of the most superficial nerve terminals imaged en face beneath the apical epithelial squames (subapical section). G) Density (count of nerve endings/field) of intraepithelial corneal nerves in cross-section at the mid-epithelial level as they run perpendicularly to the surface. H) Quantification (sum of intersections) of corneal neural complexity at the subbasal level by Sholl analysis. In F, G, and H, data are shown relative to sham-treated wt mice. I) Representative micrographs of corneal nerves from both groups at each of the three analyzed levels. All experiments were performed twice or more with 6 mice/group/experiment. For all experiments, mean±standard error of measurement is shown. To compare means, two-way ANOVA with Sidak’s posthoc test was used in B and D (treatment and time) whereas Student’s t test was applied in F, G, and H. * indicates p<0.05, ** indicates p<0.01, **** indicates p<0.0001, and ns indicates not significant.

### 6 TRPV1 signaling in the corneal tissue but not in T cells is relevant for preventing corneal neuropathy development in DED

Th1 and Th17 CD4+ T cells are pathogenic in DED as they promote apoptosis of corneal epithelial and conjunctival goblet cells and favor corneal barrier disruption(59–63). On the other hand, TRPV1 channels play a role in the activation of CD4+ T cells by enabling T cell receptor-induced calcium influx(64). Therefore, we investigated Th1 and Th17 CD4+ T cells to rule out a confounding effect of TRPV1 deficiency on their pathogenic capacity. First, we measured interferon (IFN) γ- and interleukin (IL)-17-production by flow cytometry in the CD4+ T cells of the eye-draining lymph nodes (Figures 6A-C). Wild-type and TRPV1KO mice with DED showed a non-significant trend towards more IFNγ and IL17-producing CD4+ T cells than their same-strain sham-operated littermates (2-way ANOVA: factor treatment p=0.18 and p=0.09, respectively). By contrast, the number of IFNγ- and IL-17-secreting CD4+ T cells was significantly reduced in both groups of TRPV1KO mice compared to the wild-type strain (2-way ANOVA: factor strain p<0.01 and p<0.0001, respectively). This finding is in line with the previously mentioned role of TRPV1 in potentiating CD4+ T cell activation(64). As bloodborne immune cells must gain access to the ocular surface through limbal and conjunctival vessels, we quantified conjunctival CD4+ T cells and neutrophils by flow cytometry to determine whether TRPV1 deficiency affected conjunctival immune cell recruitment. As shown in Figures 6D and 6E, there was no statistically significant difference between groups regarding the extent of CD4+ and neutrophil infiltration in the conjunctiva, but only a trend towards more cells in DED mice (2-way ANOVA: factor treatment p=0.10 and p=0.13, respectively). Altogether these results indicate that Th1 and Th17 CD4+ T cell responses are reduced in TRPV1KO mice but that this difference does not seem to be related to DED, as the number of conjunctival CD4+ T cells and neutrophils did not differ between strains.

**Figure 6.**
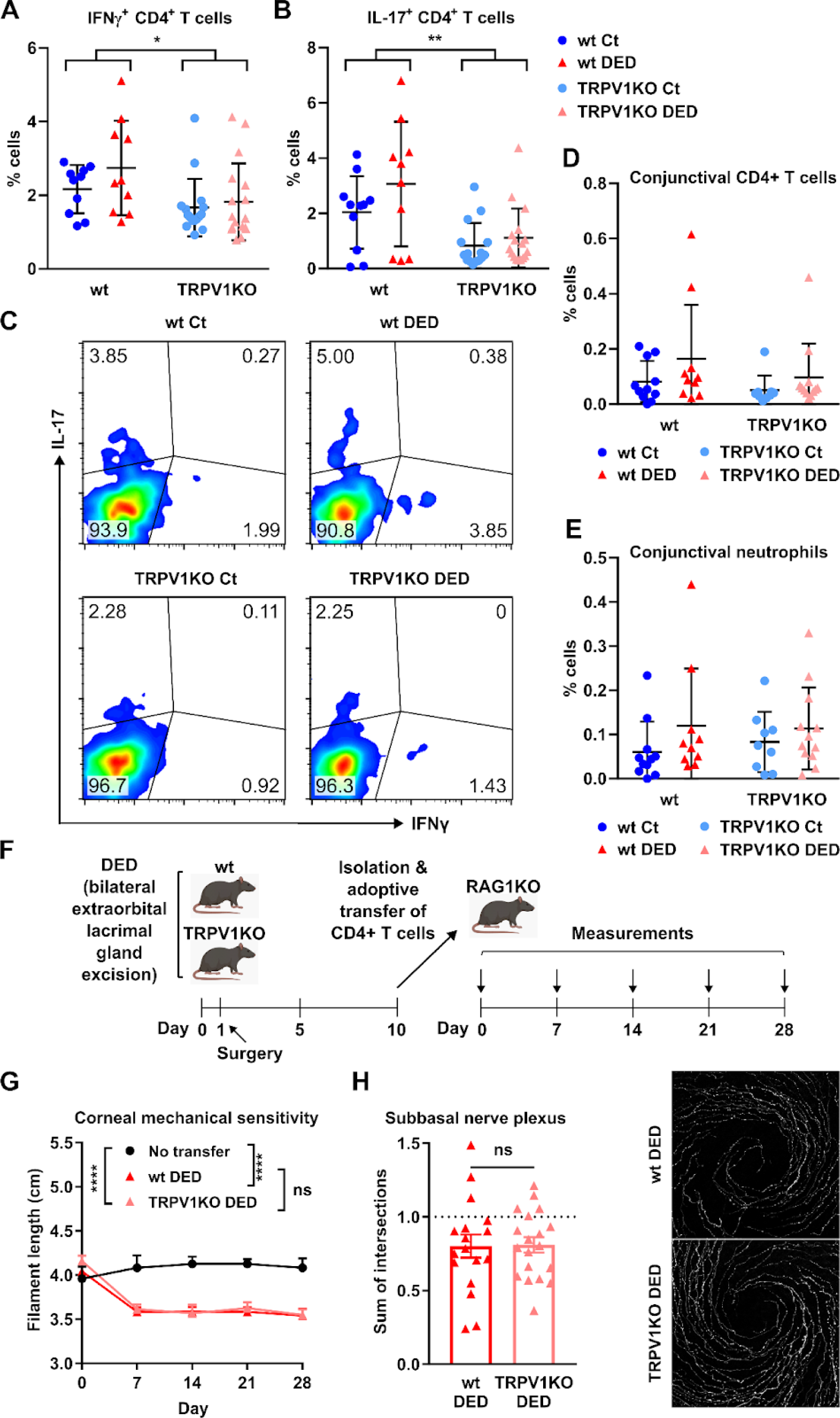
TRPV1 signaling in the corneal tissue but not in T cells is relevant for preventing corneal neuropathy development in DED. Dry eye disease (DED) was surgically induced in wild-type (wt) or transient receptor potential vanilloid-1-deficient (TRPV1KO) mice through bilateral excision of the extraorbital lacrimal gland. Sham-operated animals were included as controls (Ct). A) Interferon (IFN) γ- and B) interleukin (IL)-17-production as assessed by flow cytometry of CD4+ T cells obtained from the cervical lymph nodes of Ct and DED mice from both strains. C) Representative dot plots (IFNγ vs IL-17) of CD4+ T cells from all groups. Proportion of D) CD4+ T cells and E) neutrophils (Ly6G+) in conjunctival cell suspensions from all groups as determined by flow cytometry. F) Experimental design of adoptive transfer. DED was surgically induced in wt and TRPV1KO mice. After 10 days, CD4+ T cells were isolated from spleens and lymph nodes and 1x10^6^ cells/0.5 ml were injected intraperitoneally into each RAG1KO recipient mouse. G) Corneal mechanosensitivity as measured weekly in adoptively transferred mice. H) Quantification (sum of intersections) of corneal neural complexity at the subbasal level by Sholl analysis in corneas obtained 4 weeks after adoptive transfer. Data are shown relative to sham-treated wt mice (left) along with representative micrographs (right). All experiments were performed twice or more with 6 mice/group/experiment. For all experiments, mean±standard error of measurement is shown. To compare means, two-way ANOVA with Sidak’s posthoc test was used in A, B, D, and E (treatment and strain) and in G (treatment and time) whereas Student’s t test was applied in H. * indicates p<0.05, ** indicates p<0.01, **** indicates p<0.0001, and ns indicates not significant.

Since we have recently shown that ocular activation of Th1 CD4+ T cells independently favors corneal nerve damage(65), we reasoned that the protection from DED-induced corneal neuropathy observed in TRPV1KO mice could be due to an inhibitory effect on the neuropathogenic activity of CD4+ T cells in the ocular surface. To rule this out, we performed adoptive transfer of CD4+ T cells from either wild-type or TRPV1KO mice with DED into T cell-deficient mice (Figure 6F). We compared the pathogenic capacity of wild-type and TRPV1KO CD4+ T cells using the progression of corneal neuropathy as readout. As shown in Figure 6G, T cell-deficient mice reconstituted with CD4+ T cells from TRPV1KO mice with DED lost corneal mechanical sensitivity to the same extent as wild-type DED CD4+ T cell recipients. Consistent with the commensurable impairment of corneal mechanosensitivity, both groups of recipient mice exhibited similar changes in corneal nerve morphology. As shown in Figure 6H, the complexity of the subbasal nerve plexus (measured by the sum of intersections in Sholl analysis) decreased comparably between wild-type and TRPV1KO CD4+ T cell recipients groups (-19.9% vs -19.0%, p=0.93). Altogether these results show that the lack of TRPV1 activity in CD4+ T cells is not responsible for the phenotype observed in Figures 3 and 4. Therefore, our findings indicate that TRPV1 signaling within the corneal tissue is involved in corneal neuropathy progression in DED.

## Discussion

DED progression negatively impacts corneal nerves by an as-of-yet incompletely understood mechanism. Here we show that TRPV1 signaling is required for corneal neurodegeneration to occur in a murine model of DED. TRPV1 deficiency prevents the development of corneal nerve damage while it does not significantly affect the course of DED-induced corneal epitheliopathy. The protective effect of TRPV1 deficiency does not involve an impaired ocular surface immune response nor it stems from potentially reduced pathogenic activity of TRPV1KO CD4+ T cells, as their adoptive transfer to T cell-deficient mice leads to comparable disease as that caused by CD4+ T cells from wild-type DED mice. On the contrary, our data point towards a pathogenic role of ocular TRPV1 expression in corneal neuropathy: first, we observed sensitization to capsaicin and TRPV1-dependent pain sensation as DED progresses in our model; and second, we obtained a comparable phenotype by ocular instillation of a TRPV1 blocker. Thus, our findings suggest that TRPV1 channel overactivation in corneal nerves promotes DED-associated corneal neuropathy.

DED onset is caused by tear film instability, which brings about tear hyperosmolarity and corneal desiccation(8,9). The resulting inflammatory ocular surface setting in DED abounds in stimuli with the capacity to trigger TRPV1 signaling. TRPV1 channels allow the influx of calcium to the cell upon activation by capsaicin, heat, acidic conditions, and endogenous ligands collectively known as endovainilloids(66). Most endovainilloids with TRPV1 bioactivity are cell membrane-derived lipids that are produced during inflammation(66). Because of its polymodal nature, TRPV1 plays a key role as an integrator of inflammatory signals in nociception, i.e., the detection of damage or potentially threatening stimuli by the nervous system(67,68). TRPV1 activity is also regulated indirectly by factors that may increase (sensitize) or decrease (desensitize) the channel’s gating response to agonists(66,69). For instance, extracellular ATP released by damaged cells activates surface purinergic receptor P2Y1, triggering a cascade of intracellular events that result in TRPV1 channel phosphorylation and its subsequent sensitization(70). Many known positive and negative endogenous modulators of TRPV1 are known to be dysregulated in DED(69,71). In our model, we observed increasing sensitization to capsaicin stimulation in wild-type mice with DED, which suggests that as ocular surface inflammation worsened, the threshold for TRPV1 activation was lowered, and therefore, that TRPV1 signaling increased. Accordingly, we also recorded TRPV1-dependent spontaneous pain sensation in DED mice. Our assumption is supported by reports of enhanced corneal TRPV1 activity in other models of DED(39–43).

Contrasting the ample evidence of corneal TRPV1 overactivation taking place in DED, its consequences have not been explored in detail. We demonstrate herein that corneal TRPV1 expression is required for corneal neuropathy to appear in the context of DED, yet the underlying mechanism remains unknown. As a first possibility, we and others have previously shown that corneal TRPV1 triggering leads to the release of substance P(33,39), a proinflammatory neuropeptide that favors dendritic cell maturation and Th1 cell differentiation(72). In line with this, we recently reported that type 1 immunity in the ocular surface is sufficient to drive corneal neuropathy(65) and in this study, we observed that TRPV1KO mice with DED exhibited a trend towards decreased Th1 and Th17 immune responses (Figure 5A-B). Nonetheless, this finding does not explain the resistance to DED-induced corneal nerve damage in TRPV1KO mice because the adoptive transfer of their CD4+ T cells promotes corneal neuropathy in the recipient mice. Therefore, we believe that the protective effect must lie in corneal TRPV1 expression. As a second possibility, strong TRPV1 activation triggers the apoptosis of rat cortical neurons in vitro through a mechanism that involves extracellular calcium influx, ERK activation, and reactive oxygen species production(34). TRPV1 overstimulation has been shown to induce apoptosis in tumor cells by a similar process that involves mitochondrial instability and caspase activation(35–38). However, others have examined TRPV1-expressing neurons in the trigeminal ganglion of mice and rats with DED and found no decrease in their number(39,40). Thus, corneal neuropathy in DED is not likely to be caused by a loss of trigeminal ganglion-residing neurons that supply the eye. A third scenario relates to the interdependency of corneal epithelial cells and intraepithelial nerves(7,73). Corneal nerve endings are shed along with their ensheathing epithelial cells following a diurnal cycle(74), and on the other hand, neuronal TRPV1 activation inhibits the extension and motility of sensory neurites by regulating microtubule disassembly(75). Thus, TRPV1 overactivation might lead to corneal neuropathy in DED by interfering with the axonal growth required to replenish intraepithelial corneal nerve endings daily. Of note, corneal epithelial cell turnover increases in DED(54,55), as we confirmed in our model (Figure 1F and 1G). Finally, a fourth alternative implicates TRPV1-induced axonal degeneration. Capsaicin application in peripheral tissues causes local ablation of TRPV1-expressing nerve terminals without inducing neuronal death in the sensory ganglia(76–78). This pathway is different from capsaicin-induced neuronal apoptosis and involves the activation of calcium-dependent proteases and microtubule depolymerization in the ablated axons.

Our multilevel analysis of corneal nerve morphology sheds more light on the pathophysiology of DED-associated corneal neuropathy. In addition to comparable DED-induced corneal epitheliopathy, wild-type and TRPV1KO mice evidenced a similar extent of damage in the most superficial nerve endings. These nerve terminals are found interspersed beneath the apicalmost layer of corneal epithelial cells, hence the name subapical. The combined effects of ocular desiccation and the ensuing immune response most likely drive these changes that seem to be TRPV1-independent. However, our data also shows that the proximal segments of the corneal nerves, which are located deeper within the corneal epithelium, were spared in TRPV1KO mice but not in wild-type mice. Thus, we believe that while the most superficial (subapical) nerve endings are damaged by ocular desiccation and the immune response, TRPV1 overactivation in the same corneal nerve fibers is involved in the proximal propagation of neurodegenerative changes by any of the aforementioned mechanisms. Topical blockade of TRPV1 signaling afforded more protection to the proximal than to the distal intraepithelial corneal nerve segments in wild-type mice, confirming the distal-to-proximal progression of neural damage in DED. Our findings are in line with the more extensive reduction in the apical (distal) than in the basal (proximal) intraepithelial segments of corneal nerves reported by Stepp et al(21) using a different murine DED model. Whether the TRPV1-activating stimuli are inflammatory or tissue damage-derived remains unknown. Alternatively, TRPV1 channels are responsive to mechanical stimulation and ocular desiccation increases blinking-associated attrition of the corneal surface(79). Therefore, it is possible that deficient corneal lubrication directly contributes to TRPV1 activation in DED. More research is warranted to delineate the underlying pathogenic mechanism of TRPV1-dependent corneal nerve damage reported in this study.

Intriguingly, our data shows that TRPV1KO mice have higher corneal mechanosensitivity than wild-type mice. By contrast, others have reported that ablation of TRPV1-expressing neurons by different strategies does not affect skin mechanosensitivity(80–82). TRPV1KO mice exhibited a larger surface area of corneal subapical nerve endings yet fewer corneal mid-epithelial and subbasal nerve fibers. These findings suggest a more pronounced terminal ramification of corneal nerve endings in TRPV1KO mice, which could relate to the observed higher mechanosensitivity. Although Piezo2-expressing nerve endings were initially reported to terminate within the wing and basal epithelial cells and not reach the superficial squamous cell layer(83), a more recent study using a different methodology observed scarce Piezo2+ superficial nerve endings that also expressed TRPV1 channels(84). Of note, TRPV1 activation inhibits Piezo channel activity in other sensory neurons(85), an effect that should be absent in TRPV1KO mice. Alternatively, other channel-deficient mice exhibit dysregulated gene expression in sensory neurons(86), which might account for the altered morphology and function reported herein. Nonetheless, the expression levels of Piezo2 and transient receptor potential melastatin-8 channels in the trigeminal ganglia of sham-treated TRPV1KO and wild-type mice were comparable (Figure 4B and C). At any rate, these observations could serve as a starting point for further studies into corneal nerve biology.

Finally, one limitation of our study is that despite establishing that TRPV1 facilitates DED-associated corneal neurodegeneration, we cannot ascertain in which of the many cell types found in the cornea the non-selective cation channel plays its pathogenic role. This is because in addition to sensory nerves, corneal epithelial cells, T cells, and macrophages also express TRPV1 channels that regulate their function(64,87,88). Regarding CD4+ T cells, TRPV1 channels facilitate the activation of the Th1 effector cells that contribute to corneal nerve damage(65). However, we ruled out the potential pathogenic role of CD4+ T cell-specific TRPV1 expression in our model by adoptive transfer experiments that showed that wild-type and TRPV1KO CD4+ T cells were equally detrimental to corneal nerves. Nevertheless, our findings do not allow us to conclude whether epithelial- or macrophage-specific TRPV1 channels drive DED-induced corneal neurodegeneration. At any rate, our data indicate that corneal TRPV1 inhibition might represent a therapeutical target in DED and contributes to our understanding of how corneal neuropathy occurs in the setting of DED. At the same time, further dissection of the underlying neuropathophysiology of the disease is much needed.

## Conclusions

Our study demonstrates that ocular TRPV1 activation contributes to the development of DED-associated corneal neuropathy. We observed that ocular desiccation is sufficient to damage the most superficial nerve endings in the cornea, a process in which TRPV1 signaling is not involved. However, distal-to-proximal propagation of axonal degeneration within the corneal epithelium depends on TRPV1 signaling. Overactivation of TRPV1 channels is likely to be involved in corneal neuropathy development, as we also observed that sensitization to the TRPV1 agonist capsaicin and TRPV1-dependent pain sensation take place along with disease progression. Based on these findings, we suggest that the corneal TRPV1 pathway could be a therapeutic target for corneal neuropathy in DED and other ocular surface disorders.

## Supplementary Material

**Supplementary Video 1 - Capsaicin sensitivity testing in wild-type mice.** Representative example of capsaicin challenge (100 μM). Immediately after applying 5 μl/eye onto both eyes, the mouse is released and allowed to roam freely within an enclosed space while being video is being recorded. The number of wipes during the first 30 seconds of the sequence was counted as the response.

**Supplementary Video 2 - Capsaicin sensitivity testing in transient receptor potential vanilloid-1-deficient mice.** Representative example of capsaicin challenge (100 μM). Immediately after applying 5 μl/eye onto both eyes, the mouse is released and allowed to roam freely within an enclosed space while being video is being recorded. Fewer than 10 wipes were observed, which is comparable to the response to saline observed in wild-type mice and probably due to the non-transient receptor potential vanilloid-1-dependent discomfort elicited by the instillation of an aqueous solution onto the eye.

## Abbreviations

ANOVA: analysis of variance
DED: dry eye disease
IFN-γ: interferon-γ
IL-17: IL-17
PBS: phosphate-buffered saline
RAG1KO: recombination activating gene 1-deficient
TRPM8: transient receptor potential melastatin-8
TRPV1: transient receptor potential vanilloid-1
TRPV1KO: transient receptor potential vanilloid 1-deficient
wt: wild-type

## Declarations

### Ethics approval and consent to participate

All protocols were approved by the Institute of Experimental Medicine animal ethics committee (approval #084/2020) and adhered to the Association for Research in Vision and Ophthalmology Statement for the Use of Animals in Ophthalmic and Vision Research.

### Consent for publication

Not applicable

### Availability of data and materials

The datasets analyzed during the current study are available from the corresponding author upon reasonable request.

### Authors’ contributions

MP, AV, MG, AT, and JGG contributed to the conception and design of the work; MP, AV, AC, FS, IAK, CMS, DVA, FF, and JGG contributed to the acquisition and analysis of data; MP, AV, MG, AT, and JGG contributed to the interpretation of data; MP, AT, MG, and JGG drafted and/or substantively revised the manuscript. All authors read and approved the final manuscript.

## Acknowledgements

Not applicable

